# Establishment of a well-characterized SARS-CoV-2 lentiviral pseudovirus neutralization assay using 293T cells with stable expression of ACE2 and TMPRSS2

**DOI:** 10.1101/2020.12.26.424442

**Authors:** Sabari Nath Neerukonda, Russell Vassell, Rachel Herrup, Shufeng Liu, Tony Wang, Kazuyo Takeda, Ye Yang, Tsai-Lien Lin, Wei Wang, Carol D. Weiss

## Abstract

Pseudoviruses are useful surrogates for highly pathogenic viruses because of their safety, genetic stability, and scalability for screening assays. Many different pseudovirus platforms exist, each with different advantages and limitations. Here we report our efforts to optimize and characterize an HIV-based lentiviral pseudovirus assay for screening neutralizing antibodies for SARS-CoV-2 using a stable 293T cell line expressing human angiotensin converting enzyme 2 (ACE2) and transmembrane serine protease 2 (TMPRSS2). We assessed different target cells, established conditions that generate readouts over at least a two-log range, and confirmed consistent neutralization titers over a range of pseudovirus input. Using reference sera and plasma panels, we evaluated assay precision and showed that our neutralization titers correlate well with results reported in other assays. Overall, our lentiviral assay is relatively simple, scalable, and suitable for a variety of SARS-CoV-2 entry and neutralization screening assays.

## Introduction

In December 2019 a cluster of atypical pneumonia cases appeared in Wuhan, China. The etiological agent was later identified as severe acute respiratory syndrome coronavirus 2 (SARS-CoV-2), the causative agent of coronavirus disease 2019 (COVID-19) (1–4). In the past year, SARS-CoV-2 spread as a global pandemic with more than 75 million cases and 1.6 million deaths (Source: Johns Hopkins Coronavirus Resource Center; https://coronavirus.jhu.edu/). A key priority in fighting the ongoing pandemic involves measuring immune responses to the spike (S) glycoprotein of SARS-CoV-2, a critical target for developing preventive vaccines (5) and antibody (Ab)-based therapeutics for COVID-19 patients (6, 7), including therapeutic monoclonal antibodies (mAbs) and convalescent plasma therapy (8–16).

Assessments of serological responses to the S glycoprotein typically include virus microneutralization (MN) assays or enzyme-linked immunosorbent assay (ELISA), and ELISA variants, such as lateral flow assay (LFA), chemiluminescence immunoassay (CLIA), and electrochemiluminescence immunoassay (ECLIA) (17–19). Replicating, wild-type (WT) SARS-CoV-2 MN assays remain the gold standard, but they are labor intensive due to the need for high biosafety level containment (BSL-3) handling by trained personnel and challenges for high throughput (20). On the other hand, ELISA formats are safe and high throughput, but they do not always measure titers that strongly correlate with neutralization titers measured in the WT MN assay (10, 18, 21–23). Neutralizing Abs are thought to be an important component of protection. Some Abs that bind to S in ELISAs do not neutralize virus because they bind to S epitopes that do not interfere with receptor binding or fusion steps needed for virus entry (24–26).

The trimeric S glycoprotein mediates virus entry by binding to the ACE2 receptor on target cells and catalyzing fusion between viral and target cell membranes. Proteolytic processing of S is required for its fusion competence. The multi-basic furin-like cleavage site (RRAR*SV) allows S to be efficiently cleaved into the S1 subunit that contains the receptor binding domain (RBD) and the S2 subunit that contains domains needed for fusion (27–29). Efficient entry into the target cells additionally requires S protein priming at the S2’ site by cellular proteases, such as TMPRSS2 or cathepsins B and L (Cat B or L) (28). Depending upon the cell type, cellular proteases promote entry at the cell surface (e.g., TMPRSS2 in lung epithelium and TMPRSS4 in gut enterocytes) or in endosomes (e.g., Cat L) (28, 30). Small molecules or other inhibitors that target the S protein fusion function or cellular proteases needed for S2’ priming prevent the fusion step of entry (28). Neutralizing Abs directed against the top of the RBD typically compete with virus binding to ACE2, while those directed against the side surfaces of the RBD often do not efficiently compete with ACE2 binding and may therefore show less potent neutralization (17).

Pseudoviruses bearing viral envelope proteins provide safe surrogates for highly pathogenic viruses in MN assays. Several groups have generated SARS-CoV-2 pseudoviruses with glycoprotein defective murine leukemia virus (MLV)-, human immunodeficiency virus (HIV)-, and vesicular stomatitis virus (VSV)-based systems and used them in neutralization assays based on fluorescence (monomeric neon green) or enzymatic activity (nano-, gaussia-, and firefly luciferases) read outs in a variety of target cell types (20, 24, 31–44). In the present study, we describe our optimized conditions for an HIV-based lentiviral SARS CoV-2 pseudovirus neutralization assay. To resemble respiratory cells with TMPRSS2 and facilitate assay procedures, we established a stable 293T cell line expressing both ACE2 and TMPRSS2. We present our detailed methodology and the performance characteristics of the assay, which should be suitable for many quantitative, high-throughput virus neutralization and entry screens that can be easily performed in a routine BSL-2 laboratory.

## Materials and methods

### Ethics Statement

Use of de-identified sera and plasma samples in this study was approved the US Food and Drug Administration Research in Human Subjects Committee.

### Plasmids, cell lines and inhibitors

Full-length open reading frame of the S gene of SARS-COV2 Wuhan-Hu-1 isolate (Genbank accession: YP_009724390.1) was synthetized by GenScript (Piscataway, NJ) and cloned into the pCMV/R expression plasmid. Mutations in S were introduced using standard molecular biology protocols and confirmed by sequencing. The HIV gag/pol (pCMVΔR8.2), pCMV/R, and Luciferase reporter (pHR’CMV-Luc) plasmids described previously (45, 46) were obtained from the Vaccine Research Center (National Institutes of Health (NIH), Bethesda, MD). pCAGGS-TMPRSS2 plasmid (47) was obtained from Dr. Mikhail Matrosovich (University of Marburg, Germany). pHAGE2-EF1aInt-TMPRSS2-IRES-mCherry plasmid was obtained from Dr. Jesse Bloom (Fred Hutchinson Cancer Center, Seattle, WA) (48). pCMV-VSV-G was obtained from Dr. Kathy Bouir, (University of California, San Diego). A plasmid encoding human ACE2 (hACE2-TM) was obtained from the NIH Vaccine Research Center. HEK293T-ACE2 (293T.ACE2_s_) cells stably expressing ACE2 were obtained through BEI Resources, National Institute of Allergy and Infectious Diseases, NIH; (NR-52511, contributed by Jesse Bloom, Fred Hutchinson Cancer Research Center, Seattle, WA) (32). The 293T, Vero, Vero E6, A549, Caco-2, Calu-3 and Huh-7 cells were maintained at 37°C in Dulbecco’s modified eagle medium (DMEM) supplemented with high glucose, L-Glutamine, minimal essential media (MEM) non-essential amino acids, penicillin/streptomycin and 10% fetal bovine serum (FBS). Chemical inhibitors, camostat mesylate (TMPRSS2 inhibitor; Cat no: SML0057) and chloroquine (endosomal acidification inhibitor; Cat no: 50-63-5) were obtained from MilliporeSigma.

### Antibodies and sera

Mouse mAb 10G6H5 against SARS-COV2 S protein was purchased from GenScript (Piscataway, NJ). Rabbit antisera against the S1 subunit, the receptor binding domain (RBD), and the S2 subunit of SARS-COV2 S protein (49) were provided by Surender Khurana (US Food and Drug Administration, Silver Spring, MD). The National Institute of Biological Standards and Control (NIBSC) serology reference panel, including 20/120, 20/122, 20/124, 20/126, 20/128 and 20/130, against SARS-COV-2 was provided by Giada Mattiuzzo (National Institute for Biological Standards and Control (NIBSC), Potters Bar, UK). Human plasma from COVID-19 patients were provided by James Rost and Norton Elson (Washington Adventist Medical Healthcare), Nicholas Cacciabeve (Advanced Pathology Associates), and Rob San Luis, Hollie Genser, Demetra Collier, Meaza Belay, Genevieve Caoili, Zanetta E. Morrow, and Bruana Streets (Quest Diagnostics). A serum and plasma proficiency panel (focused concordance samples) with high, medium, and low neutralizing titers against SARS-COV-2 and a blinded serum and plasma panel developed for the SARS-CoV-2 neutralization assay concordance survey (SNACS) were provided by Dr. David Montefiori (Duke University, Durham, NC). Negative control sera collected in September-December of 2009 from 45 volunteers ages 48-64 years was described previously (50). HIV-1 p24 hybridoma (183-H12-5C) was obtained through the AIDS Research and Reference Reagent Program, Division of AIDS, NIAID, NIH and contributed by Dr. Bruce Chesebro.

### Pseudovirus production and neutralization

Pseudoviruses bearing the S glycoprotein and carrying a firefly luciferase (FLuc) reporter gene were produced in 293T cells. Briefly, 5μg of pCMVΔR8.2, 5μg of pHR’CMVLuc and 0.5μg of S or its mutants (codon optimized) expression plasmids with or without 2μg of the TMPRSS2 expression plasmid were co-transfected in 293T cells. Pseudovirus supernatants were collected approximately 48 h post-transfection, filtered through a 0.45μm low protein binding filter, and used immediately or stored at −80°C. Pseudovirus titers were measured by infecting different cells for 48 h prior to measuring luciferase activity (luciferase assay reagent, Promega, Madison, WI), as described previously (51). Pseudovirus titers were expressed as relative luminescence unit per milliliter of pseudovirus supernatants (RLU/ml).

Neutralization assays were performed on 293T cells transiently transfected or transduced with ACE2 and TMPRSS2 genes for stable expression. Briefly, pseudoviruses with titers of approximately 10^6^ RLU/ml of luciferase activity were incubated with antibodies or sera for one hour at 37°C. Pseudovirus and antibody mixtures (100 μl) were then inoculated onto 96-well plates that were seeded with 3.0 x 10^4^ cells/well one day prior to infection. Pseudovirus infectivity was scored 48 h later for luciferase activity. The antibody dilution or mAb concentration causing a 50% and 80% reduction of RLU compared to control (ID_50_ and ID_80_ or IC_50_ and IC_80_, respectively) were reported as the neutralizing antibody titers. Titers were calculated using a nonlinear regression curve fit (GraphPad Prism software Inc., La Jolla, CA). The mean 50% and 80% reduction of RLU compared to control from at least two independent experiments was reported as the final titer. For experiments involving camostat mesylate (0.03-500 μM) and chloroquine (0.39-25 μM), target cells were treated with each inhibitor for two hours before pseudovirus infection in the presence of respective inhibitor.

### Generation of transient and stable 293T-ACE2.TMPRSS2 cells

The 293T-ACE2.TMPRSS2_t_ cells transiently expressing low, medium, and high levels of TMPRSS2 were generated by co-transfection of ACE2-TM and pCAGGS-TMPRSS2 plasmids in 2μg, 4μg and 8μg, respectively. Cell surface expression levels of ACE2 and TMPRSS2 were determined by flow cytometry. Briefly, transfected cells were harvested using non-enzymatic cell dissociation solution (Sigma-Aldrich) and resuspended in flow cytometry staining buffer (FCSB), PBS containing 2% FBS and 0.1% sodium azide, at 10^7^ cells/ml. Cells were incubated in AF647-conjugated anti-ACE2 mAb (Santa Cruz: sc-390851) and/or AF488 conjugated anti-TMPRSS2 mAb (Santa Cruz: sc-515727), followed by three FCSB washes. Cells were then fixed in 2% formaldehyde, washed with FCSB and quantified for signal intensity using a BD LSRFortessa-X20 flow cytometer (Becton Dickinson).

To generate stable 293T-ACE2.TMPRSS2_s_ cells, VSV-G-pseudotyped lentiviruses carrying the human TMPRSS2 gene were generated by co-transfecting 293T cells with the pHAGE2-EF1aInt-TMPRSS2-IRES-mCherry, pCMVΔ8.2, and pCMV-VSV-G plasmids. Packaged lentivirus was used to transduce 293T-ACE2 cells in the presence of 10μg/mL polybrene, and the resulting bulk transduced population was single-cell sorted into clear bottomed 96 well plates by flow cytometry that was based on intermediate or high mCherry positivity on a BD FACSAria II Cell Sorter. Once single cell clones reached confluence, they were screened for mCherry/TMPRSS2 expression via EVOS Floid cell imaging station (Thermo Fisher, Waltham, MA), and several clones with visible mCherry expression were expanded. For verifying mCherry expression via flow cytometry, cells were harvested with enzyme-free cell dissociation buffer (Thermo Fisher, Waltham, MA), washed, and resuspended in FACS buffer. One clone that displayed intermediate levels of mCherry expression and maximum pseudovirus infectivity titer was selected and referred to as 293T-ACE2.TMPRSS2_s_. Up to the present, this clone has supported high-level infectivity of SARS-CoV-2 pseudoviruses through 20 passages.

### SARS-CoV-2 mNG infection and confocal microscopy

Vero E6 cells, 293T-hACE2_s_, and 293T-ACE2.TMPRSS2_s_ cells were seeded on poly-L-lysine-coated coverslips one day prior to infection. Infection with live SARS-CoV-2-mNG (MOI:0.1) was carried out in medium containing 2% FBS for one hour at 37°C, prior to washing the cells twice with PBS and then maintaining in culture, described above. SARS-CoV-2-mNG expressing mNeon Green (mNG) in place of ORF7 was described previously (52). At 24 h post infection (p.i.) SARS-CoV-2-mNG-infected coverslips were fixed with 4% paraformaldehyde in PBS at room temperature for 20 min followed by PBS washes. SARS-CoV-2-mNG infection and fixing procedures were performed in a BSL-3 laboratory at the US Food and Drug Administration. Coverslips were counterstained with Hoechst 33258 dye (Thermo Scientific) and mounted on microscope slides with Fluoromount-G (SouthernBiotec). Confocal microscopy was performed by using SP8 DMI6000 confocal microscope (Leica Microsystems Inc, Germany) equipped with 25x water immersion objective lens and 405, 488 and 594 laser lines for Hoechst, mNG and mCherry signal, respectively.

### Immunoblot analysis

Pseudoviruses were resolved by SDS-PAGE and detected by Western blot using a mouse mAb (183-H12-5C) against HIV-1 p24 Gag and rabbit antisera against the S1 subunit and the S2 subunit of SARS-COV-2 S protein.

### Statistics analysis

To evaluate assay precision, six NIBSC plasma standards, 15 focused concordance samples and 21 SNACS samples were tested by three operators. Two operators ran four independent experiments (two independent experiments per operator), and a third operator ran one experiment. Titers were calculated from curves using eight dilutions. Intermediate precision, expressed as the percent coefficient of variation (%CV), was assessed separately for ID_50_ and ID_80_ titers. Sample dilutions with observed titers of less than 1:40 were considered as negative for antibodies to SARS-CoV-2 and were imputed a value of 1:20. An exploratory analysis was additionally performed by excluding titers of less than 1:40. Samples with more than 50% of less than 1:40 were excluded from all analyses. The total %CV accounts for both inter-operator and inter-assay variability and were estimated as follows based on a linear mixed model of the natural log-transformed titers with sample as a fixed effect and operator as a random effect:

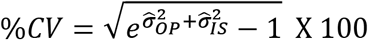

where 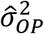 and 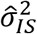 were the estimated inter-operator and inter-assay (within-operator) variance components from the model, respectively. *SAS version 9.4* was used to perform the linear mixed model analysis.

To evaluate accuracy, since the true titers of test samples are not known, the Spearman correlation coefficient between the reported titers and the “observed” titers was estimated with GraphPad Prism software.

## Results and discussion

### Selection of optimal target cells for SARS-CoV-2 pseudovirus infectivity

We generated SARS-CoV-2 pseudoviruses bearing full-length S glycoprotein from the SARS-COV-2 Wuhan-Hu-1 isolate using a second-generation lentiviral packaging system that we used previously for producing pseudoviruses bearing other viral glycoproteins (51, 53, 54). We assessed SARS-CoV-2 pseudoviruses for infectivity in target cell types that were previously reported to support various levels of pseudovirus and replicating virus infectivity. A previous study reported higher infectivity of VSV-based pseudoviruses in Huh7, 293T, and Vero cells compared to CHO, MDCK, and HepG2 cells (39), but other studies reported poor or no infectivity of 293T cells due to the absence of ACE2 receptor (33, 43). Other cell types, including stably engineered cells (293T, 293T17, HT1080, BHK21), transiently transfected cells (293T, Caco-2, Huh7, HepG2, MDCK), and various continuous cell lines (Vero-E6, A549, BEAS2B, Calu-3, H1299, MRC5, Caco-2, HeLa, K562) that express ACE2, TMPRSS2, or both, have been widely reported to support pseudovirus infectivity to different degrees (20, 24, 31–44). We therefore assessed a panel of cell types, including Vero, Vero E6, A549, Caco-2, Calu-3, Huh7, 293T, and 293T cells transiently expressing ACE2 (293T-ACE2t), TMPRSS2 (293T-TMPRSS2_t_), or both (293T-ACE2.TMPRSS2_t_), to identify the cells that supported the highest levels of infectivity for our pseudoviruses.

As expected, our SARS-CoV-2 pseudoviruses lacked infectivity above background levels (approximately 10^4^ RLU/ml) in 293T cells due to the absence of the ACE2 receptor, while Vero-E6 cells are naturally resistant to human lentivirus infection due to intrinsic restriction factors (Fig 1A). The A549, Caco-2, Calu-3, and Huh-7 cells also lacked infectivity above background levels (Fig 1A). Transient 293T-TMPRSS2_t_ or 293T-ACE2t cells had 5.7- and 40-fold signal above background, respectively, while transient 293T-ACE2.TMPRSS2_t_ cells resulted in a 3130-fold signal above background (Fig 1A). A stable 293T-ACE2_s_ cell line displayed 144-fold higher signal compared to background (Fig 1A).

**Figure 1.**
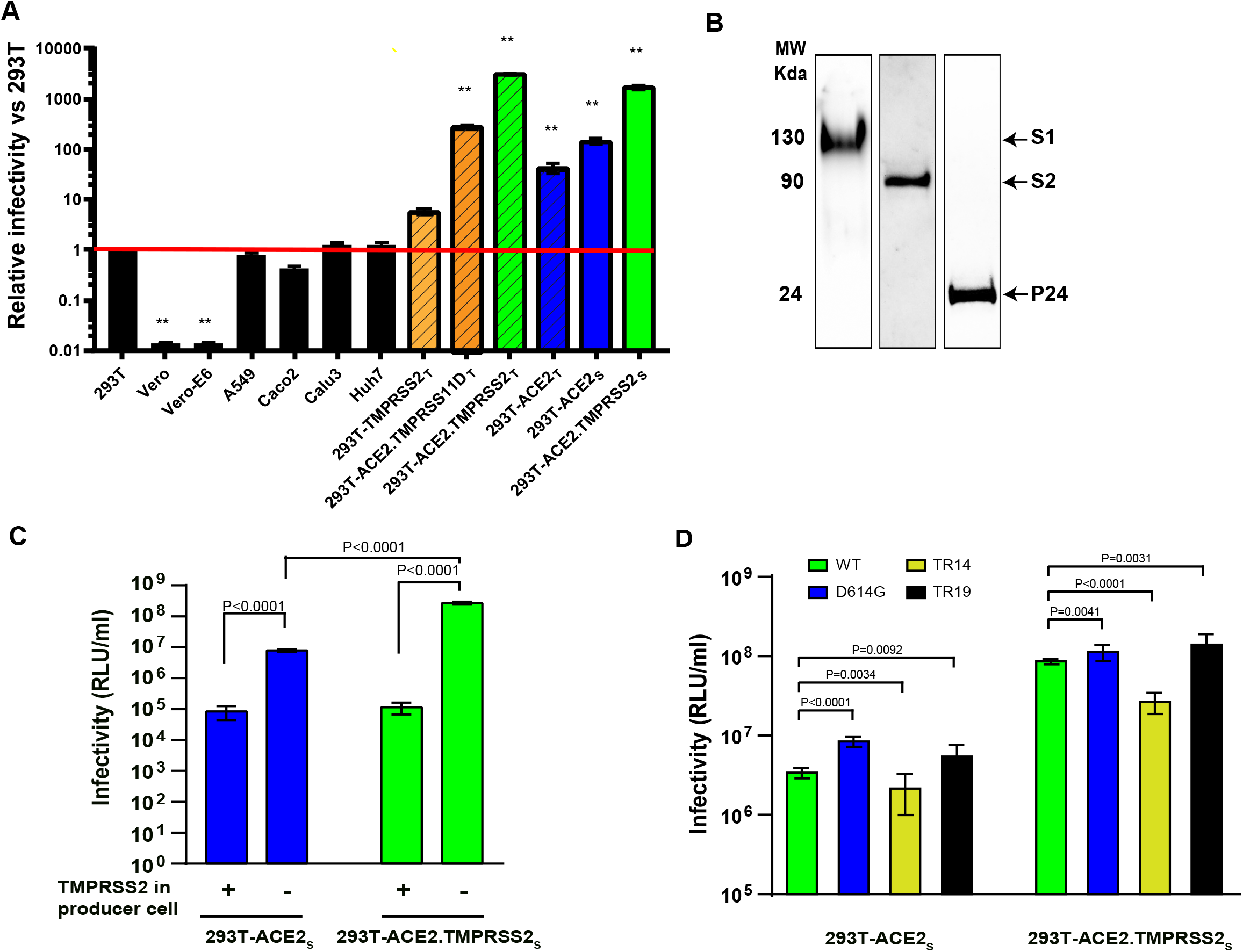
SARS-CoV-2 lentiviral pseudovirus infectivity under various conditions. (A) Relative infectivity of SARS-CoV-2 pseudoviruses in various target cells. The Y-axis shows relative infectivity compared to background in 293T cells. Subscript ‘t’ or ‘s’ refers to transient or stable expression, respectively. Red line indicates background level (1 x 10^4^ relative units of luciferase activity/ml). (B) Western blots probed for spike S1/S2 subunits and HIV p24 incorporated into pseudoviruses. (C) Infectivity on 293T-ACE2_s_ and 293T-ACE2.TMPRSS2_s_ of pseudoviruses primed with or without TMPRSS2 during pseudovirus production. (D) Infectivity on 293T-ACE2_s_ and 293T-ACE2.TMPRSS2_s_ of pseudoviruses bearing full-length, wildtype S glycoprotein (WT), an S glycoprotein with the D614G substitution, or S glycoproteins with C-terminal truncations of 14 (TR14) and 19 (TR19) amino acids. Data are shown as means and standard deviations from 3-4 independent experiments (panel A, C and D). The tests for two-group comparison were analyzed using GraphPad Prism software. P values of 0.05 were considered statistically significant. **: P<0.0001, compared to the infectivity in 293T.

A prior report demonstrated priming of SARS-CoV-2 S in VSV-based pseudoviruses by TMPRSS2-related proteases TMPRSS11D, 11E, 11F, and 13 in Calu-3 target cells. Of the TMPRSS2-related proteases, TMPRSS13 was as efficient as TMPRSS2, while the others were less efficient in promoting S priming and thus infectivity (41). In the present study, we investigated the ability of TMPRSS11D (also known as human airway trypsin) in target 293T cells transiently expressing ACE2 and TMPRSS11D (293T-ACE2.TMPRSS11Dt). Consistent with the prior report, TMPRSS11D was less efficient in S priming, as reflected by an 11-fold lower infectivity in 293T-ACE2.TMPRSS11Dt cells compared to 293T-ACE2.TMPRSS2_t_ cells (Fig 1A).

To facilitate SARS-CoV-2 pseudovirus neutralization assays and partly mimic natural SARS-CoV-2 target cells that express TMPRSS2, we established a stable cell line expressing both ACE2 and TMPRSS2 (293T-ACE2.TMPRSS2_S_) by transducing the 293T-ACE2 cells with a lentivirus encoding TMPRSS2 and mCherry as a bicistrionic transcript. The ACE2.TMPRSS2_S_ cells conferred infectivity approximately 1700-fold above background, confirming the contribution of TMPRSS2 protease activity for priming the S protein for fusion competence. Because the ACE2.TMPRSS2_S_ cells facilitated greater levels of infectivity and provided > 10^8^ RLU/ml infectivity (>3 log range) for resolving titers, we focused on qualifying the 293T-ACE2.TMPRSS2_s_ cells for our future studies. We also confirmed S protein incorporation into pseudoviruses and proteolytic processing of the S protein to generate S1 and S2 subunits that migrate at 130kDa and 90kDa, respectively (Fig 1B).

### Optimization of SARS-CoV-2 pseudovirus infectivity

We investigated several conditions for optimizing SARS-CoV-2 pseudovirus production in 293T cells. When comparing S priming by TMPRSS2 during pseudovirus production to priming during entry into target cells, we found that co-expressing TMPRSS2 during pseudovirus production reduced pseudovirus infectivity, possibly due to TMPRSS2-induced premature activation of S that promotes conformational changes to fusion-incompetent or post-fusion structures (Fig 1C). This finding is consistent with a previous report suggesting the importance of tight regulation of protease cleavage of the S protein for preserving SARS-CoV-2 infectivity (55).

We also investigated variant S proteins to further optimize pseudovirus production. We generated S genes with the D614G mutation or a C-terminal cytoplasmic tail truncation of 19 amino acids (TR19) that were previously reported to yield higher infectivity titers compared to full length WT S glycoprotein (34, 38, 39, 42, 56). The D614G change represents a natural mutation outside RBD that became dominant in the circulating strains (39). In 293T-ACE2_s_ cells, the D614G mutation was reported to confer 1-log or >1/2-log higher infectivity to VSV-based or lentivirus-based S pseudoviruses, respectively (38, 39, 57). Furthermore, viruses with the G614 S were associated with higher virion stability and increased *in vitro* SARS-CoV-2 replication fitness in primary human airway epithelial cells and Calu-3 cells (57). Increased infectivity conferred by the D614G change is due to removal of hydrogen-bond interaction with T859 from a neighboring protomer of the S trimer. This results in an allosteric change of RDB domain to an “up” conformation that facilitates ACE2 receptor binding that may make the virus modestly more susceptible to neutralization by some sera or antibodies, depending the epitopes targeted by the antibodies (39, 56, 57). Truncation of C-terminal 18 or 19 amino acids, which removes a putative ER retention signal, was also demonstrated to enhance HIV-based pseudotyping efficiency by 10-fold compared to WT S protein in 293T-ACE2_s_ cells (42, 58). The higher infectivity conferred by the C-terminal cytoplasmic tail truncation of 19 amino acids may be due to higher number of infectious particles (42).

Consistent with previous reports, we found that pseudoviruses bearing G614 and TR19 S proteins displayed 0.5- and 0.2-log higher infectivity, respectively, compared to WT pseudovirus in 293T-ACE2_s_ cells (Fig 1D). Pseudoviruses with TR14 S had slightly lower infectivity compared to WT pseudoviruses (Fig 1D). However, in 293T-ACE2.TMPRSS2_s_ cells the pseudovirus bearing the G614 S and TR19 S were more similar to WT S pseudoviruses, but the pseudovirus bearing the TR14 S displayed 0.5 log lower infectivity compared to WT pseudovirus (Fig 1D). Infectivity titers of all pseudoviruses were 1-1.5-log lower on 293T-ACE2_s_ cells compared to 293T-ACE2.TMPRSS2_s_ cells (Fig 1D). Based on these studies, we used WT S to qualify our pseudovirus neutralization assay because it represents the native, full-length spike, and the infectivity of WT S pseudoviruses give a large dynamic range for generating neutralization dose-response curves. Additional efforts to enhance pseudovirus infectivity with polybrene, a polycation that is known to enhance lentiviral transduction efficiency by minimizing charge-repulsion between the virus and cells, displayed no effect on pseudovirus infectivity.

### Replication of SARS-CoV-2-mNG in 293T-ACE2.TMPRSS2_s_ cells

We confirmed the acceptability of the 293T-ACE2.TMPRSS2_s_ cells for SARS-CoV-2 infectivity and neutralization studies by assessing how well the cells support infection by replicating SARS-CoV-2. We compared replication levels and cytopathic effects (CPE) in 293T-ACE2.TMPRSS2_s_ to Vero E6 cells that are widely used for the propagation of SARS-CoV-2, as well as to 293T-ACEs cells that lack TMPRSS2. Typically, SARS-CoV-2-induces CPE in Vero cells by 48-72 h post infection (p.i.), characterized by cell rounding, detachment, degeneration, and syncytia (59). However, by 24 h p.i., when both SARS-CoV-2-infected Vero E6 and 293T-ACE2_s_ cells began to display mNG expression and early CPE, SARS-CoV-2-infected 293T-ACE2.TMPRSS2_s_ cells displayed robust mNG expression and higher levels of CPE with nearly 50% of infected monolayer undergoing detachment (Fig 2). The rapid kinetics of infection is consistent with a preprint indicating co-expression of ACE2 and TMPRSS2 synergistically increases SARS-CoV-2 or pseudovirus entry efficiency (40).

**Figure 2.**
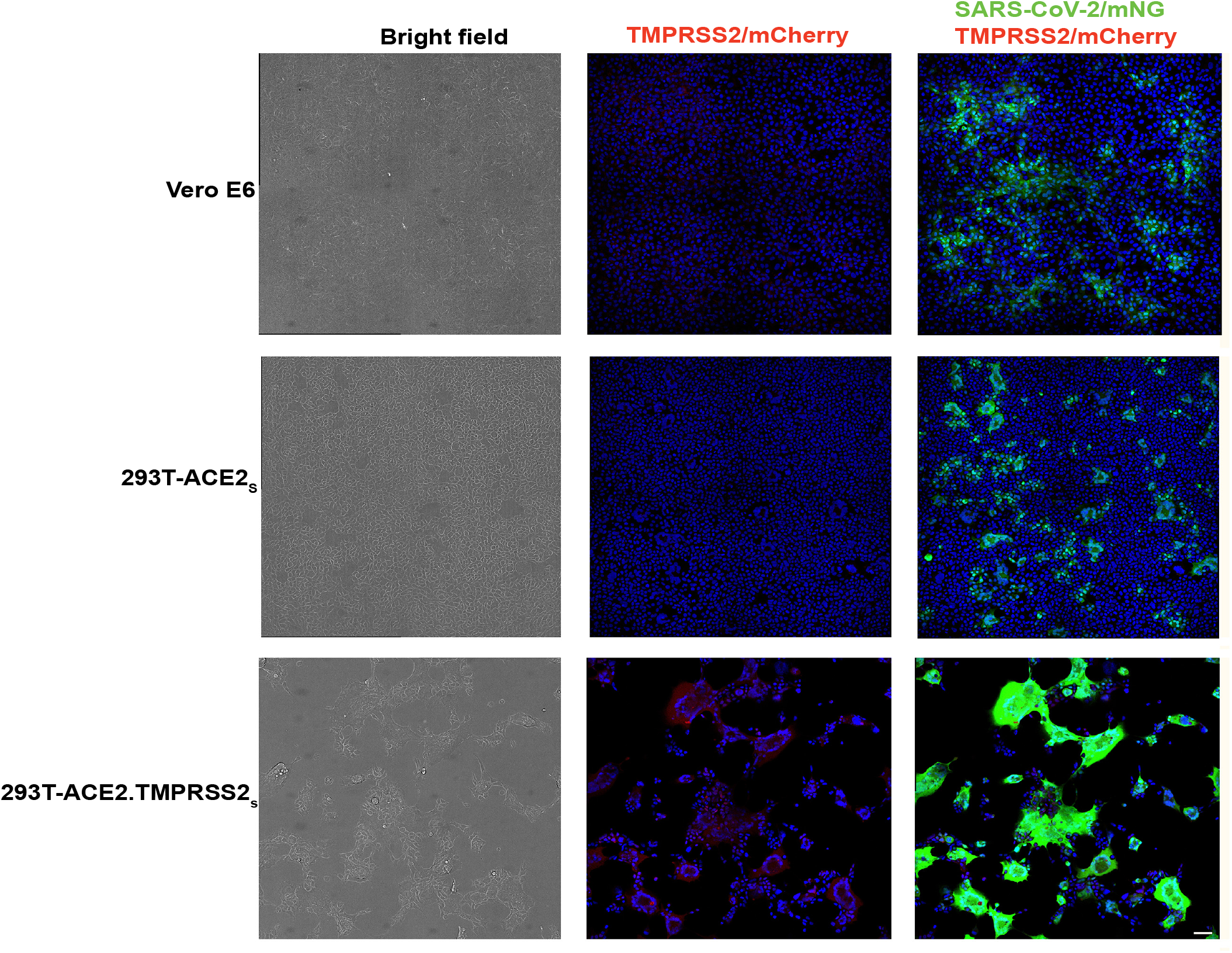
Infection of Vero E6, 293T-ACE2_s_ and 293T-ACE2.TMPRSS2_s_ with replicating SARS-CoV-2-mNG virus. Cells inoculated with 100 PFU/ml of virus were fixed and imaged at 24 hours post infection by confocal microscopy. The left column shows bright field images (black and white), with the 293-ACE2.TMPRSS2 cells (bottom) showing a high degree of cytopathic effect and syncytium formation, resulting in fewer cells. The center column shows merged images of TMPRSS2/mCherry and Hoechst dye (blue) with only the 293-ACE2.TMPRSS2 cells (bottom) showing positive signals in red. The right column shows merged images of SARS-CoV-2/mNG (green), TMPRSS2/mCherry (red), and Hoechst dye (blue). Compared to Vero (top) and 293-ACE2 (middle), the 293T-ACE2.TMPRSS2 (bottom) cells show stronger SARS-CoV-2 nNG signals in green. Scale indicates 50um.

### Entry pathways of SARS-CoV-2 pseudoviruses

Next, we used chemical inhibitors to determine the protease that primes SARS-CoV-2 S protein for membrane fusion during pseudovirus entry in 293T-ACE2.TMPRSS2_s_ cells. SARS-CoV-2 can use pH-dependent and -independent pathways for cell entry (60). In cells lacking TMPRSS2, SARS-CoV-2 relies on endosomal-pH-dependent cysteine protease, such as cathepsin L for S priming, while entry is predominantly dependent on priming by TMPRSS2 in natural airway cells, such as lung epithelial cells, type II pneumocytes (60, 61). We found that pseudovirus entry into 293T-ACE2_S_ cells, which lack TMPRSS2, was sensitive to the endosomal pH acidification inhibitor chloroquine, with a half maximal inhibitory concentration [IC_50_] of 0.79μM, but relatively insensitive to a TMPRSS2 inhibitor camostat mesylate. In contrast, pseudovirus entry was sensitive to camostat mesylate [IC_50_: 0.88μM] in 293T-ACE2.TMPRSS2_s_ cells, but much less so to chloroquine treatment (Fig 3). Thus, SARS-CoV-2 predominantly uses TMPRSS2 for priming S during virus entry into the 293T-ACE2.TMPRSS2_s_ cells.

**Figure 3.**
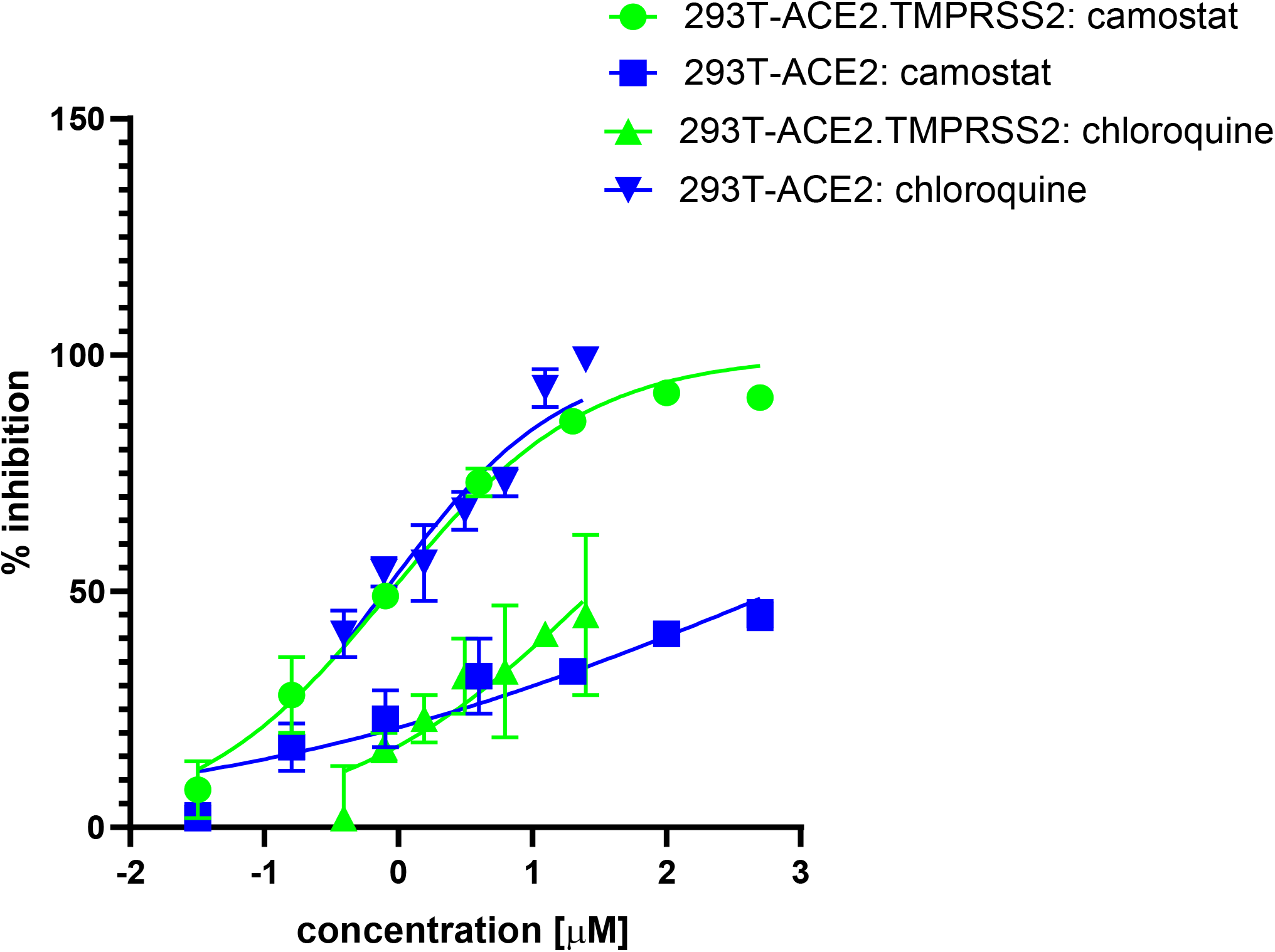
pH-dependent and -independent pathways of cell entry of SARS-CoV-2 S pseudoviruses. Camostat mesylate inhibits TMPRSS2 activity and chloroquine inhibits endosomal acidification required for cathepsin activity. Cells were pretreated with camostat mesylate or chloroquine for 2 h prior to pseudovirus infection in the presence of inhibitor for a period of 48 h. Results shown are representative of three independent experiments.

### Optimization of SARS-CoV-2 pseudovirus inoculum for neutralization assays

We next determined the inoculum range that would assure consistent neutralization titers according to the law of mass action (62). Serial dilutions of rabbit serum or mAb (10G6H5) were mixed with four different pseudovirus inoculums over a three-log range, prior to incubation with 293T-ACE2.TMPRSS2_S_ cells (Fig 4). Although 100% neutralization was achieved at high serum or mAb concentrations using a relatively low inoculum of 2 x 10^4^ relative luciferase units/ml (RLU/ml), the dose-response curve displayed high variation at higher dilutions, precluding generation of a reliable curve for calculating 50% neutralization titers (Fig 4A and B). However, inoculums in the 4 x 10^5^ – 1.4 x 10^7^ RLU/ml range generated overlapping curves with little variation (Fig 4C and D). These dose-response curves yielded 50% neutralization titers with serum inhibitory dilution (ID_50_) and mAb inhibitory concentration (IC_50_) values that varied less than 2-fold over all pseudovirus inoculums. Therefore, inoculums of 5 x 10^5^ – 1 x 10^7^ RLU/ml were used for the neutralization assay.

**Figure 4.**
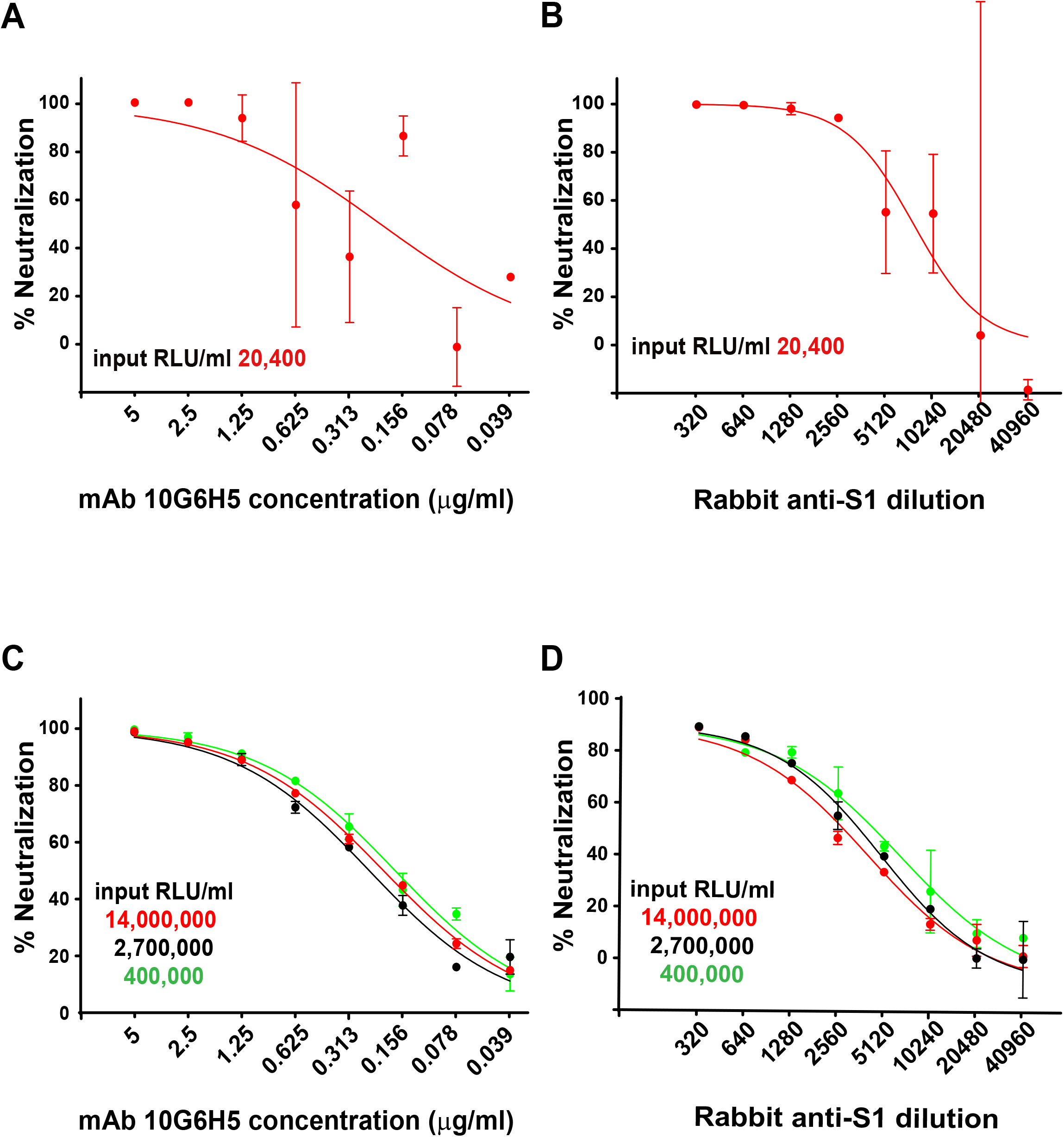
Optimization of pseudovirus inoculum for neutralization assays. Neutralizations were performed in 293T-ACE2.TMPRSS2_s_ cells with mAb 10G6H5 (A and C) and a rabbit serum against the S1 subunit (B and D) using various pseudovirus inoculums.

### Assessment of the influence of ACE and TMPRSS2 levels on neutralization titers

Because the 293T-ACE2.TMPRSS2_s_ cells may have higher levels of TMPRSS2, ACE2, or both, compared to some primary airway cells, we explored whether different levels of ACE2 and TMPRSS2 on target cells might influence neutralization titers. We transiently transfected 293T cells with ACE2 and TMPRSS2 to achieve low, medium, and high levels of ACE2 and TMPRSS2. Expression levels of ACE2 and TMPRSS2 were confirmed via flow cytometry (Fig 5). Transfection of a higher plasmid concentration of ACE2 and TMPRSS2 resulted in an increase in the number of ACE2^+^/TMPRSS2^+^ cells (Fig 5A) as well as cell surface expression (Fig 5B and 5C). Neutralization assays performed with rabbit sera against RBD or S1 subunit, murine mAb 10G6H5, as well as an NIBSC reference plasma (#20/130), showed no significant differences in neutralization titers among the target cells with different levels of ACE2 and TMPRSS2 (Table 1). Neutralization ID_50_ titers of rabbit sera against RBD and S1 subunit ranged from 9377 to 10540 and 5462 to 6742, respectively. NIBSC reference plasma ID_50_ titers ranged from 2355 to 3130, while negative control sera lacked neutralization activity. IC_50_ values for the mAb 10G6H5 ranged from 0.119 to 0.197μg/ml. The 80% neutralization titers (ID_80_ or IC_80_) were also similar among target cells with different levels of ACE2 or TMPRSS2. These findings indicate that levels of ACE2 and TMPRSS2 may not have a significant impact on neutralization titers for many antibodies.

**Figure 5.**
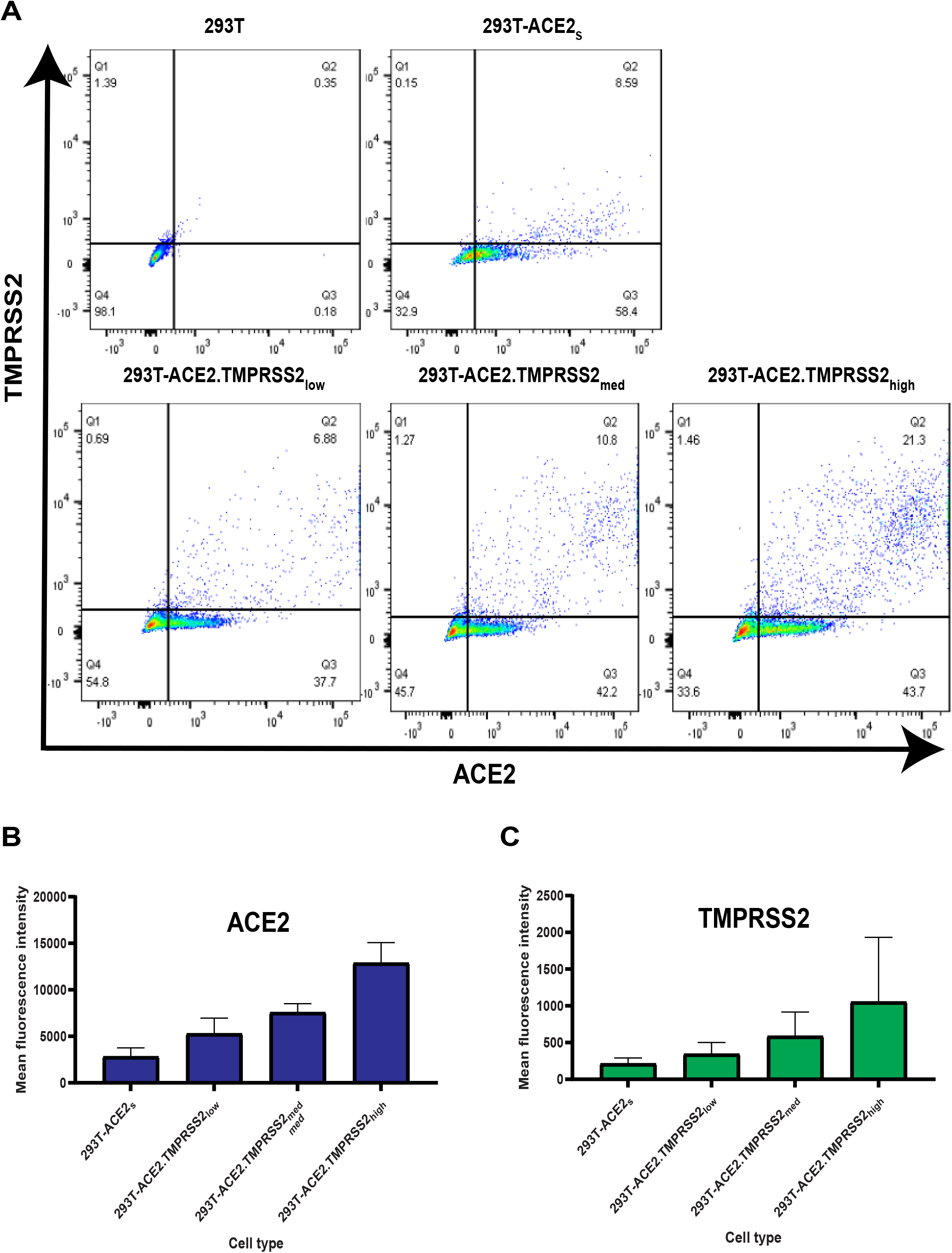
ACE2 and TMPRSS2 levels at the cell surface. (A) Cell surface expression of ACE2 and TMPRSS2 on various cell types. (B) Mean fluorescence intensity (MFI) of ACE2 on various cell types compared to 293T cells (C) Mean fluorescence intensity (MFI) of TMPRSS2 on various cell types compared to 293T cells

**Table 1:**
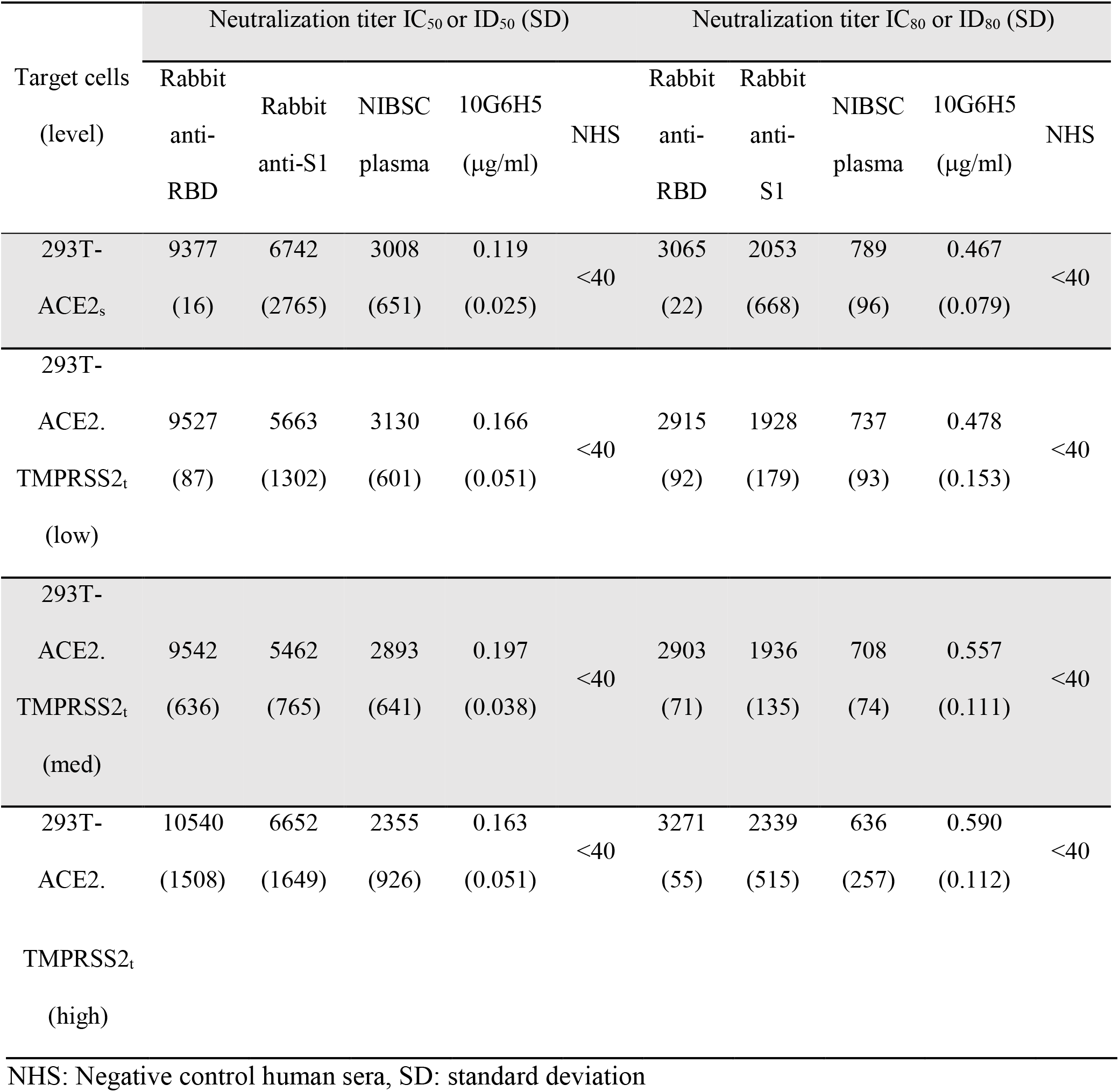
Neutralization titers in 293T cells with low, medium, and high levels of ACE2 and TMPRSS2

### Assessment of neutralization specificity and range of antibody titers

We assessed assay specificity and range of antibody titers using sera with reported neutralization titers, as well as 15 plasma samples from patients hospitalized with COVID-19. Thirty sera collected before 2019 served as negative controls, along with a negative control reference plasma standard (NIBSC #20/126) (Fig 6). Positive controls included the focused concordance samples comprising four high, five medium, and five low neutralizing antibody titers. All negative control sera failed to neutralize SARS-CoV-2 pseudoviruses at the lowest dilution tested (1:40). Neutralization titers (ID_50_ and ID_80_) segregated into high, medium, and low groupings, consistent with reported titers (Fig 6A and 6B). Plasma for patients hospitalized with acute COVID-19 showed a wide range of titers, consistent with previous reports (63). We note, however, that the presence of reverse transcriptase or integrase inhibitors in sera or plasma samples from persons on anti-retroviral therapy (ART) has the potential to interfere with lentiviral pseudovirus readout that is dependent on reverse transcription and integration of the reporter gene. We therefore use lentiviral pseudoviruses bearing an envelope protein from amphotropic MLV or VSV as an additional control when assessing clinical samples that could include subjects on therapeutic or preventive ART. Non-specific inhibition of MLV- or VSV-pseudoviruses identifies sera that cannot be evaluated using this assay.

**Figure 6.**
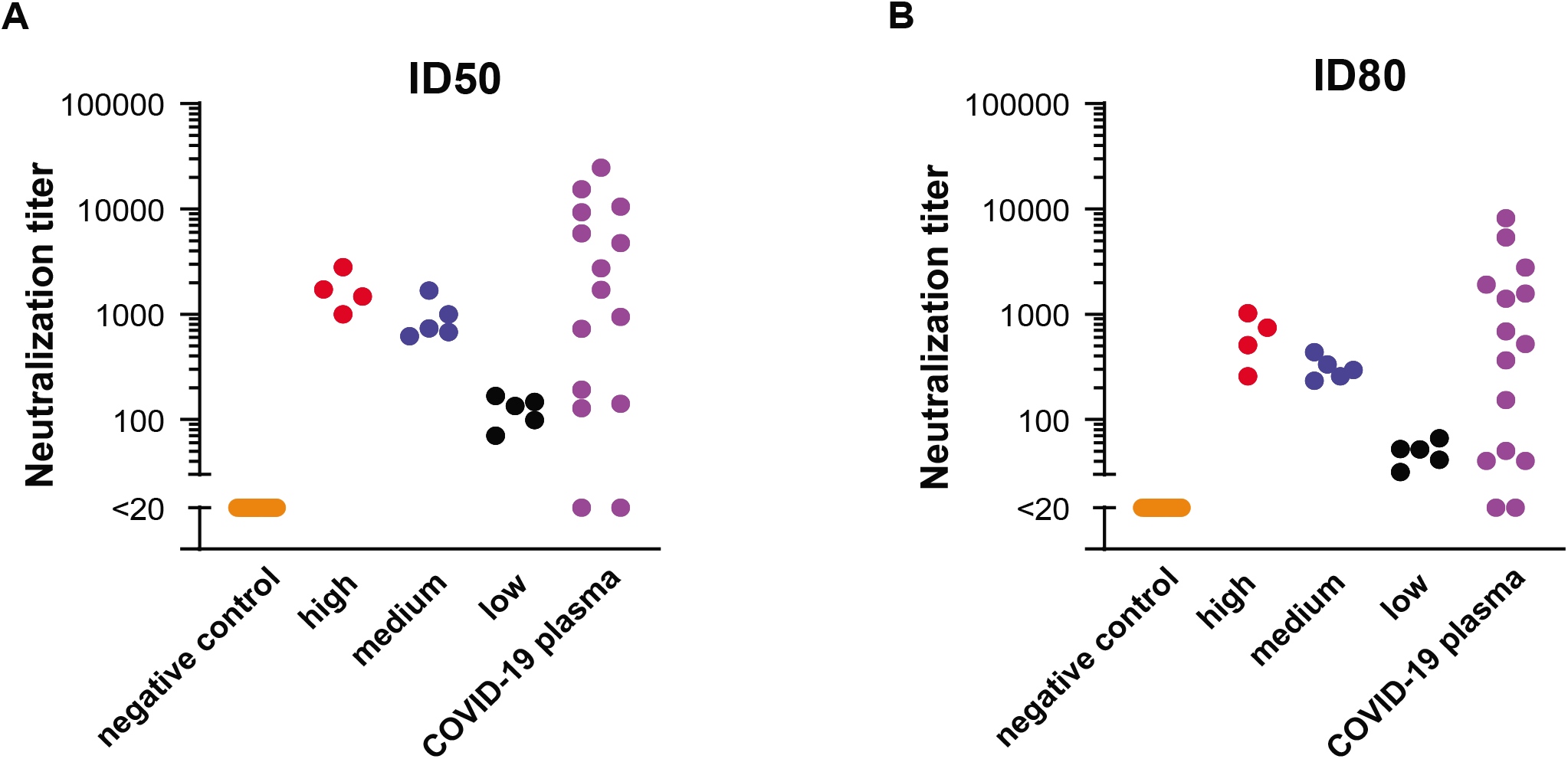
Neutralization titers of serum panels. ID_50_ (A) and ID_80_ (B) titers. ID_50_ = inhibitory dilution leading to 50% neutralization compared to control. ID_80_ = inhibitory dilution leading to 80% neutralization compared to control.

### Assessment of assay precision

To further qualify the assay performance, we assessed intermediate precision among three operators using blinded test samples that included the six NIBSC plasma standards, 15 sera samples from the focused concordance samples panel, and a blinded panel of 21 sera samples that was used in a survey to assess assay concordance among labs using various SARS-CoV-2 neutralization assays (https://dhvi.duke.edu/duke-team-implement-sars-cov-2-neutralization-assay-concordance-survey-laboratories-worldwide). Neutralization titers giving 50% or 80% inhibition (ID_50_ and ID_80_, respectively) compared to control were used to calculate the %CV. Only positive samples (with at least 50% of titer results ≥ 1:40) were included in the precision calculation, which excluded six samples for ID_50_ and 13 samples for ID_80_.The overall %CV across all samples for ID_50_ and ID_80_ titers was 38.8% and 30.8%, respectively, when titers < 1:40 were imputed to be 1:20. We consider these results to be acceptable for a neutralization assay and adequate for most clinical studies. When titers <1:40 were excluded from the analysis, the %CV was 27.5% and 20.7%, respectively.

### Assessment of inter-laboratory agreement

We used the samples with reported neutralization titers as a benchmark for assessing accuracy of our assay. Although the reported titers were generated using different neutralization assay formats, we nevertheless found a strong correlation between our titers and the neutralization titers reported for the focused concordance samples (Fig 7A and B) and several NIBSC reference standards (Fig 7C and D). These results provide assurance that our assay provides titers that correlate well with titers measured in other assay formats.

**Figure 7.**
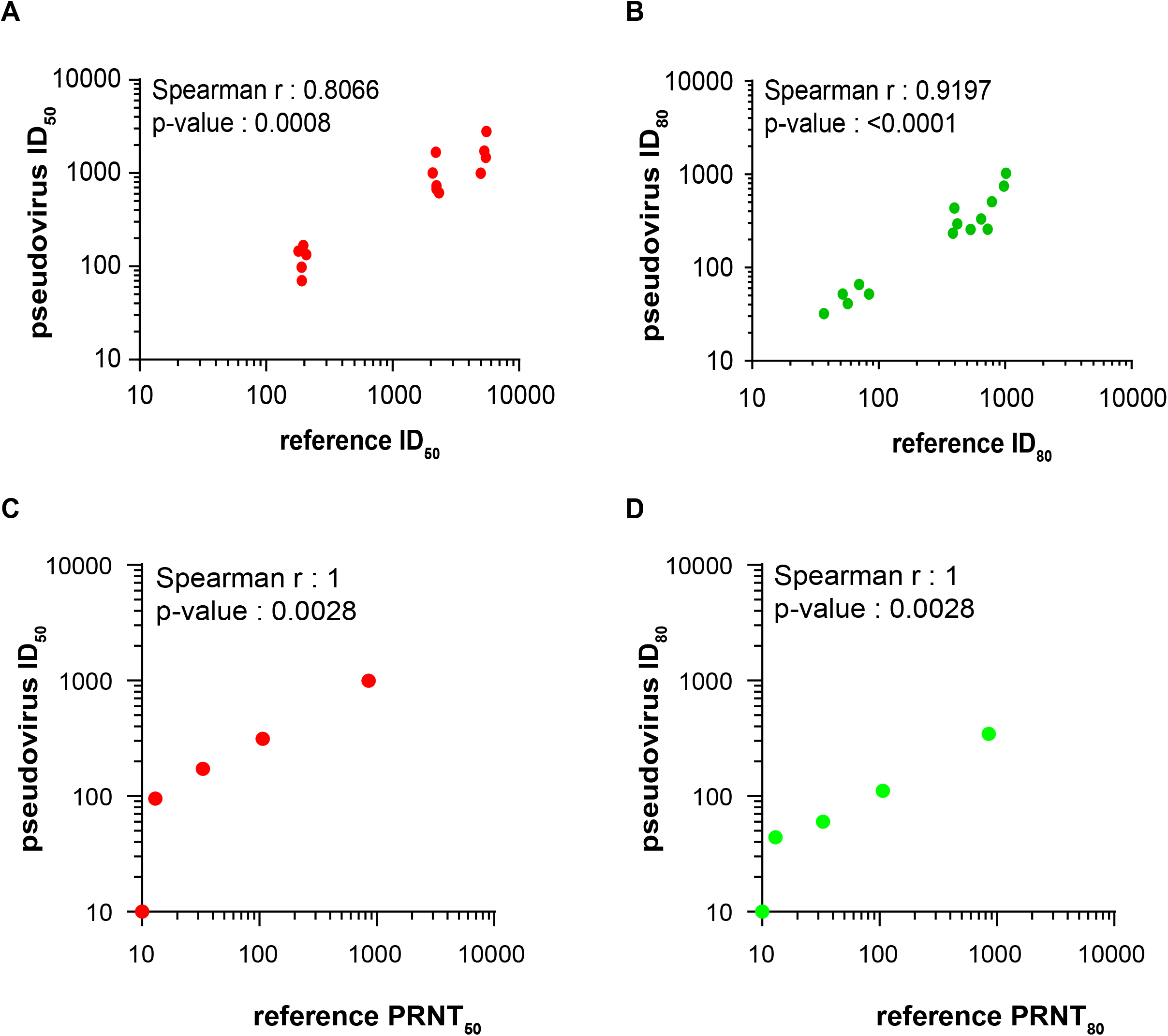
Correlation of neutralization titers between different neutralization assays. Spearman correlation of ID_50_ (A and C) or ID_80_ (B and D) values comparing titers generated in the present study to (A and B) the focused concordance samples and (C and D) an NIBSC reference standards with reported titers generated by a plaque reduction neutralization (PRNT) assay.

## Conclusion

We describe optimized procedures and detailed performance characteristics of an HIV-based, lentiviral pseudovirus neutralization assay for SARS-CoV-2 using a stable 293T cell line expressing ACE2 and TMPRSS2. The assay is quantitative, has a large dynamic range, and generates titers that correlate well with titers generated in other assays. The safety and relative simplicity of the assay makes it a valuable and versatile tool for evaluating mAb potency and neutralizing antibody titers in a BSL-2 lab setting.

## Acknowledgments

We thank the following for generously contributing materials for this study: Dr. David Montefiori (Duke University), the HIV Vaccine Trials Network, and the HIV Prevention Trials Network for the focused concordance samples; Dr. Michael Busch (Vitalant Research Institute) for the SNACS samples; Jesse Bloom and Katharine H. D. Crawford (Fred Hutchinson Cancer Center) for the 293T-ACE2 cells and plasmids. We also thank Dr. Carolyn A. Wilson (US Food and Drug Administration) for facilitating receipt of plasma samples from subjects hospitalized for COVID-19 and Drs. Konstantin Virnik and Ira Berkower (US Food and Drug Administration) for critical reading of the manuscript.

